# Machine Learning Models for Segmentation and Classification of Cyanobacterial Cells

**DOI:** 10.1101/2024.12.11.628068

**Authors:** Clair A. Huffine, Zachary L. Maas, Anton Avramov, Chris Brininger, Jeffrey C. Cameron, Jian Wei Tay

## Abstract

Timelapse microscopy has recently been employed to study the metabolism and physiology of cyanobacteria at the single-cell level. However, the identification of individual cells in brightfield images remains a significant challenge. Traditional intensity-based segmentation algorithms perform poorly when identifying individual cells in dense colonies due to a lack of contrast between neighboring cells. Here, we describe a newly developed software package called Cypose which uses machine learning (ML) models to solve two specific tasks: segmentation of individual cyanobacterial cells, and classification of cellular phenotypes. The segmentation models are based on the Cellpose framework, while classification is performed using a convolutional neural network named Cyclass. To our knowledge, these are the first developed ML-based models for cyanobacteria segmentation and classification. When compared to other methods, our segmentation models showed improved performance and were able to segment cells with varied morphological phenotypes, as well as differentiate between live and lysed cells. We also found that our models were robust to imaging artifacts, such as dust and cell debris. Additionally, the classification model was able to identify different cellular phenotypes using only images as input. Together, these models improve cell segmentation accuracy and enable high-throughput analysis of dense cyanobacterial colonies and filamentous cyanobacteria.

## Introduction

Timelapse microscopy, combined with the use of fluorescent labeling and sensing, allows molecular processes to be observed in individual cells at a sub-cellular level. Recent techniques have shown that cyanobacteria, a model organism for the study of photosynthetic processes, can be filmed over extended periods of time^1–3^. The resolution of microscopy datasets has led to discoveries that were not previously observed in bulk culture experiments^4,5^ such as the regulation of photosynthetic processes^2^ and organelle development and positioning^6–8^.

Due to the large number of individual cells that can be captured in a single image, computational pipelines are often used to obtain single-cell data from microscopy datasets. However, the identification (or segmentation) of individual cells in the resulting images remains the main bottleneck in these pipelines. Cell segmentation is typically performed using intensity-thresholding, where every pixel above a set threshold is identified as being part of a cell^3,9,10^. Intensity-thresholding is popular as it is a relatively simple technique that works well if cells are fluorescently labeled, so that the fluorescence signal is much brighter compared to the background.

Cyanobacteria produce photosynthetic pigments which are autoflourescent. While this might initially seem advantageous, the fluorescence signal is typically non-uniform throughout the cell. Additionally, the fluorescence intensity changes depending on the cell’s photosynthetic capacity, which can lead to issues when choosing a threshold intensity. Together, these issues have meant that using photosynthetic fluorescence to identify individual cells is undesirable. Alternatively, fluorescent proteins or dyes could be used to mark the cells. However, the presence of the autofluorescence once again complicates matters since most microscopes are limited to imaging ∼3 fluorescence channels due to spectral overlap^11,12^. Requiring a fluorescent label would limit one’s ability to label other molecules or organelles of interest. It is therefore advantageous to develop segmentation algorithms which use the brightfield image, which is generated by light transmitted through the sample, to remove the need for further labeling.

We previously developed intensity-thresholding algorithms to identify cyanobacteria in brightfield images. However, these images are difficult to segment because there is little contrast between the cell interior and the background^13^. The problem is exacerbated when cells grow in dense colonies or for filamentous strains of cyanobacteria as cell boundaries become even less pronounced (Fig. 1, S1).

**Fig 1:**
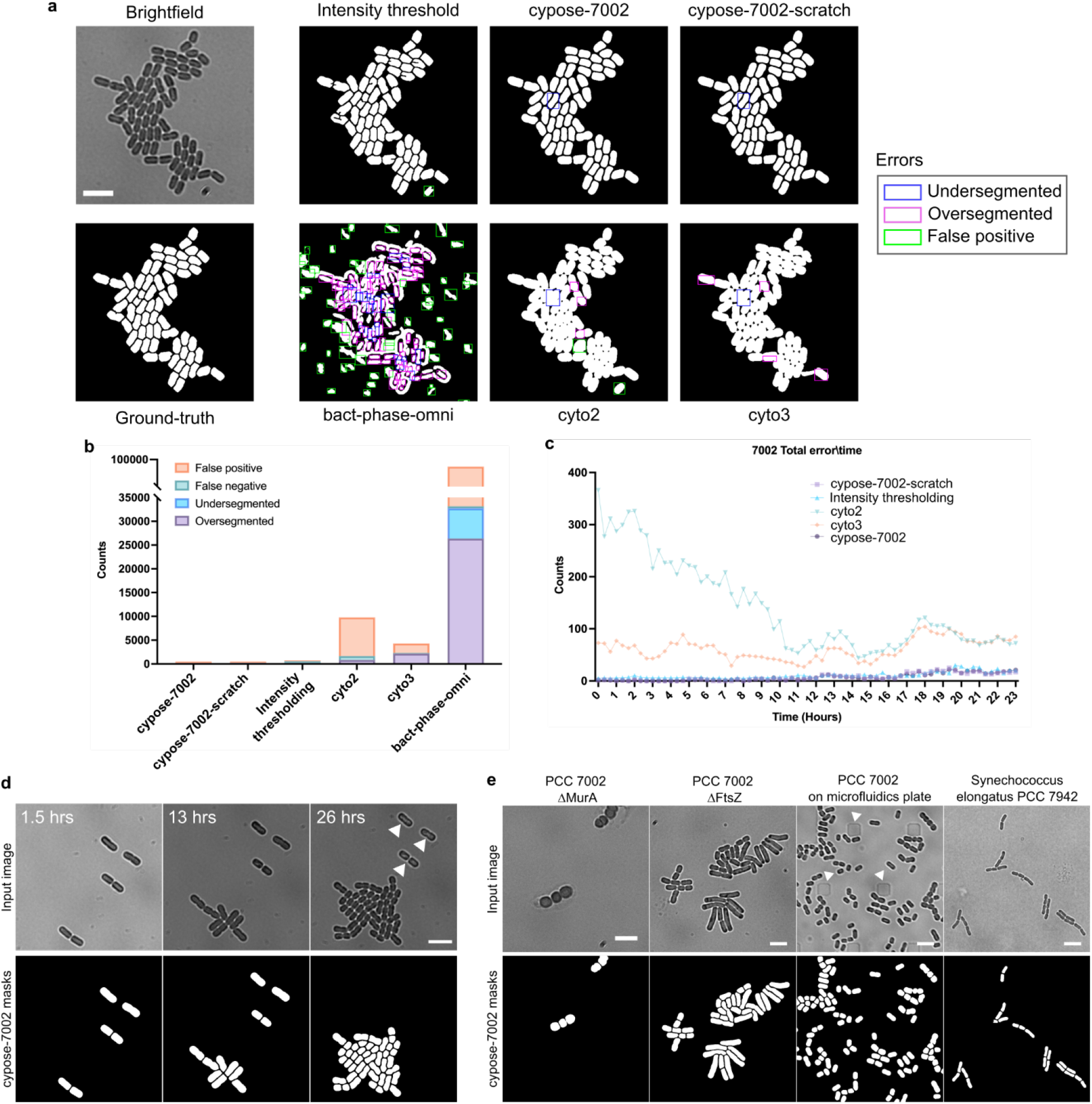
(a) Representative images of PCC 7002 showing the input image, ground-truth, and resulting masks from the intensity-thresholding algorithm and the various machine learning models. Errors in each mask are highlighted by a box: undersegmentation in blue, oversegmentation in magenta, and false positives (additional objects) in green. Scale bar indicates 5 µm. (b) Bar plot showing the total number of segmentation errors over the entire benchmark movie. (c) Plot showing the number of total errors at each frame. The bact-phase-omni model gave a total error over two orders of magnitude higher and is excluded for clarity (data shown in Fig. S2). (d) Representative frames from a movie showing that the cypose-7002 model was able to identify and ignore dead cells (indicated by white arrows), which were identified by their lack of growth during the movie. Scale bar indicate 5 µm. (e) Representative frames showing the model’s ability to identify different species/phenotypes and under different imaging conditions. The white arrows indicate posts of the microfluidic plate. Scale bars indicate 5 µm.

Here, we describe the development of a family of machine learning models, collectively named Cypose, to improve the segmentation of individual cyanobacterial cells. Compared to intensity-based thresholding, machine learning segmentation models can learn to identify cells using complex hierarchical image features and have been shown to work well even without fluorescent labeling^14,15^. Additionally, we describe a method to train a classification model, using a convolutional neural network (CNN) named Cyclass, to perform image-based identification of different cellular phenotypes. We demonstrate that the Cyclass model can be used to identify different cellular phenotypes using only image data as input. We demonstrate the usefulness of both Cypose and Cyclass by showing that these models can be used together to initially segment cyanobacterial cells in dense colonies, then classify different cellular phenotypes in a timelapse video. This methodology could be helpful in studies of mixed bacterial species by enabling multiple genotypes/phenotypes to be imaged simultaneously or to distinguish individual species in studies containing mixed populations.

## Results

### Development of the Cypose cyanobacterial segmentation models

Our Cypose segmentation models are fine-tuned models based on the Cellpose base models^16^. Cellpose consists of a U-Net like convolutional neural network, which transforms an image into a series of spatial gradients. These gradients are then used to identify and label individual cells in an image.

In initial testing, we found that the base Cellpose cytoplasmic models (cyto2^17^ and cyto3^18^) showed poor performance when segmenting brightfield images of cyanobacteria (Fig. 1a). This is likely because these models were trained on cytoplasmic images of eukaryotic cells^16^, which have different phenotypic and morphological features compared to cyanobacteria. We also tested a separate segmentation model, bact-phase-omni, from the Omnipose package^19^ (which itself is derived from Cellpose), which was trained on images of bacteria. However, we found that this model appeared to perform worse than Cellpose for segmentation, likely because it was trained primarily on phase contrast images.

To achieve higher quality segmentation on cyanobacteria, we trained a family of specialist models. Three different models were trained: (1) cypose-7002 and (2) cypose-7002-scratch were trained on images of *Synechococcus sp*. PCC 7002 (hereafter PCC 7002), which are unicellular, while (3) cypose-33047 was trained on images of *Anabaena sp*. ATCC 33047 (hereafter ATCC 33047), which are filamentous. Both cypose-7002 and cypose-33047 were fine-tuned from the Cellpose cyto2 base model, while cypose-7002-scratch was trained from scratch on the Cellpose architecture. Details of the training and datasets used are provided in the methods section below. We note that our models are based on the Cellpose 2.0 cyto2 model rather than the recently released Cellpose 3.0 cyto3 model, as the latter was unavailable at the start of this work. However, we have included comparisons of our new models with cyto3.

### Segmentation of *Synechococcus* PCC 7002 cells

We used the cypose-7002 model to segment timelapse videos of PCC 7002 capturing single cells developing into colonies. A representative image showing a dense colony from a late frame taken from the benchmark movie is shown in Fig. 1a. Masks generated by our cypose-7002 model, the Cellpose and Omnipose models, as well as our previously described intensity-thresholding algorithm are shown for comparison. We note that while the intensity-thresholding algorithm performed well, segmentation errors tended to develop within individual cells, as well as between cells after 3 doublings (S1). However, our new Cypose model performed well over long timelapse movies. The model begins to fail when the colonies grow so dense that the cells start to overlap.

To assess the accuracy of our model, we compared the mask generated by each method to our ground-truth masks pixel-by-pixel by calculating the Intersection over Union (IoU) score, as well as the typical precision and recall scores for each method, as shown in Table 1 and S2. These scores showed that cypose-7002 outperforms other models across nearly all metrics. The cypose-7002 model appears to perform slightly worse compared to the intensity-based approach. However, as detailed in the methods, the benchmark dataset was generated using the intensity-based approach so there is likely a bias towards this approach.

**Table 1.** Performance of segmentation models calculated by comparing the generated masks with ground-truth masks pixel-by-pixel or by identifying errors in individual objects.

| Organism | Model | Pixel-based performance |  |  | Object-based performance |  |  |  |  |
| --- | --- | --- | --- | --- | --- | --- | --- | --- | --- |
|  |  | IoU | Precision | Recall | Over-segmented | Under-segmented | False Negative | False Positive | Total |
| PCC 7002 | cypose-7002 | 0.929 | 0.953 | 0.973 | 59 | 153 | 2 | 274 | 488 |
|  | cypose-scratch | 0.921 | 0.944 | 0.974 | 43 | 191 | 10 | 256 | 500 |
|  | cyto3 | 0.784 | 0.949 | 0.818 | 2175 | 12 | 128 | 1985 | 4300 |
|  | cyto2 | 0.597 | 0.853 | 0.666 | 834 | 18 | 829 | 8107 | 9788 |
|  | bact-phase-omni | 0.120 | 0.437 | 0.142 | 26364 | 6374 | 432 | 58982 | 92152 |
|  | Intensity thresholding | 0.951 | 0.966 | 0.984 | 33 | 175 | 389 | 151 | 748 |
| ATCC 33047 | cypose-33047 | 0.875 | 0.849 | 0.757 | 629 | 934 | 81 | 140 | 1784 |
|  | cyto3 | 0.732 | 0.905 | 0.793 | 33 | 2201 | 5597 | 46 | 7877 |
|  | cyto2 | 0.642 | 0.774 | 0.790 | 72 | 1199 | 2325 | 703 | 4299 |
|  | bact-phase-omni | 0.040 | 0.374 | 0.043 | 164 | 809 | 688 | 1485 | 3146 |
|  | Intensity thresholding | 0.659 | 0.798 | 0.790 | 26605 | 2373 | 904 | 727798 | 757680 |

We note that while these pixel-based metrics are typically reported when comparing cell segmentation models, they do not accurately capture inaccuracies, such as over- and under-segmentation, in the resulting objects. For example, we observed that single cells were often split into two distinct objects by both cyto2, cyto3, bact-phase-omni. In this case, the error could be as small as a single-pixel wide line, which means that the number of erroneous pixels were much smaller compared to the number of pixels in both cells, the precision and recall scores appear high.

To obtain a more accurate analysis, we developed code to recognize errors in the identified objects. We identified four errors: (1) oversegmentation (2) undersegmentation, (3) false positives, and (4) false negatives. We found that the cypose-7002 model resulted in a much lower number of errors compared to all other models, including the intensity-based approach (Table 1, Fig. 1b and b, S2). On the benchmark movie, cypose-7002 had 10% of the total number of segmentation errors compared to the other tested models. Additionally, we found that cypose-7002 only generated two false negatives, compared to the hundreds generated by the other models. We note that while the number of errors in a frame increases slightly over the duration of the movie as colonies become denser (Fig. 1c), this model remains more accurate than the other tested models.

During creation of the training dataset, lysed cells were not annotated. Consequently, we found that our cypose-7002 model was able to differentiate living and lysed cells (Fig. 1d). Additionally, to test the breadth of the model’s capability to segment cells with different morphologies, we applied the model to images of *Synechococcus elongatus* sp. PCC 7942 (hereafter PCC 7942), two genetic mutants of PCC 7002 with markedly different phenotypes, as well as images of PCC 7002 growing in a microfluidic device (Fig. 1e). In the latter, plastic posts in the microfluidic chamber are visible in brightfield. In all cases, we found that our new model correctly segmented cells, while ignoring other artifacts such as the posts.

### Comparison of scratch-trained and fine-tuned models

As previously mentioned, our cypose-7002 model was fine-tuned from the pretrained cyto2 model. The cyto2 model was trained on a diverse training set, primarily comprising of images of eukaryotic cells, with additional non-cell images containing repeating patterns, such as shells and rocks. Since no images of cyanobacteria existed in the cyto2 dataset, we wanted to test if a segmentation model trained from scratch only on cyanobacterial images would perform better compared to the fine-tuned cypose-7002 model.

The cypose-7002-scratch model was trained from scratch using the Cellpose architecture. This model was trained on 3.5x more cell images compared to cypose-7002. To account for different cell morphologies, our dataset included various PCC 7002 mutants which showed morphological deviations from WT cells (e.g., cell swelling and elongation) similar to those shown in Fig. 1e (note that these images were not shown to the fine-tuned cypose-7002 model). Results from this segmentation model are shown in Fig. 1a. Benchmark metrics were also calculated and are shown in Table 1.

Overall, we found that the scratch-trained model provided very similar results to the fine-tuned cypose-7002 model. However, training from scratch was both labor intensive (since more training data needed to be manually curated) and required substantially more training than fine-tuning. Considering the similarity in performance, we concluded that training from scratch did not offer notable advantages.

### Segmentation of filamentous cyanobacterial strains

We fine-tuned a second model (cypose-33047) to segment filamentous *Anabaena sp*. ATCC 33047. This strain is challenging to segment as it forms interconnected structures with minimal intensity differences between neighboring cells. Additionally, ATCC 33047 differentiates into three morphologically and phenotypically distinct cell types: photosynthetic vegetative cells, specialized nitrogen-fixing heterocysts, which have no or little photosynthetic pigments^20^, and akinetes^21^, which are large, spore-like cells formed during low nutrient conditions. To increase the distinction between neighboring cells and to account for the different cell types, the cypose-33047 model was trained on images of both the brightfield and the chlorophyll fluorescence channels.

The resulting masks are shown in Fig. 2a. We found that our fine-tuned model provided significant improvement when segmenting filamentous cyanobacteria compared to the Cellpose and Omnipose models, and to the intensity-thresholding method (Table 1, Fig. 2b and c, S4, S5). As before, we used a timelapse movie with cells starting from small filaments as a benchmark. To capture a variety of conditions, we selected three distinct temporal subsets of this movie for testing, capturing variable cell densities at the start, middle, and end. We found that the cypose-33047 model excelled in early and mid-movie frames, with cyto3 performing better in later frames. This suggests that our model could be improved by increasing the number of images showing dense filaments in our training dataset.

**Fig 2:**
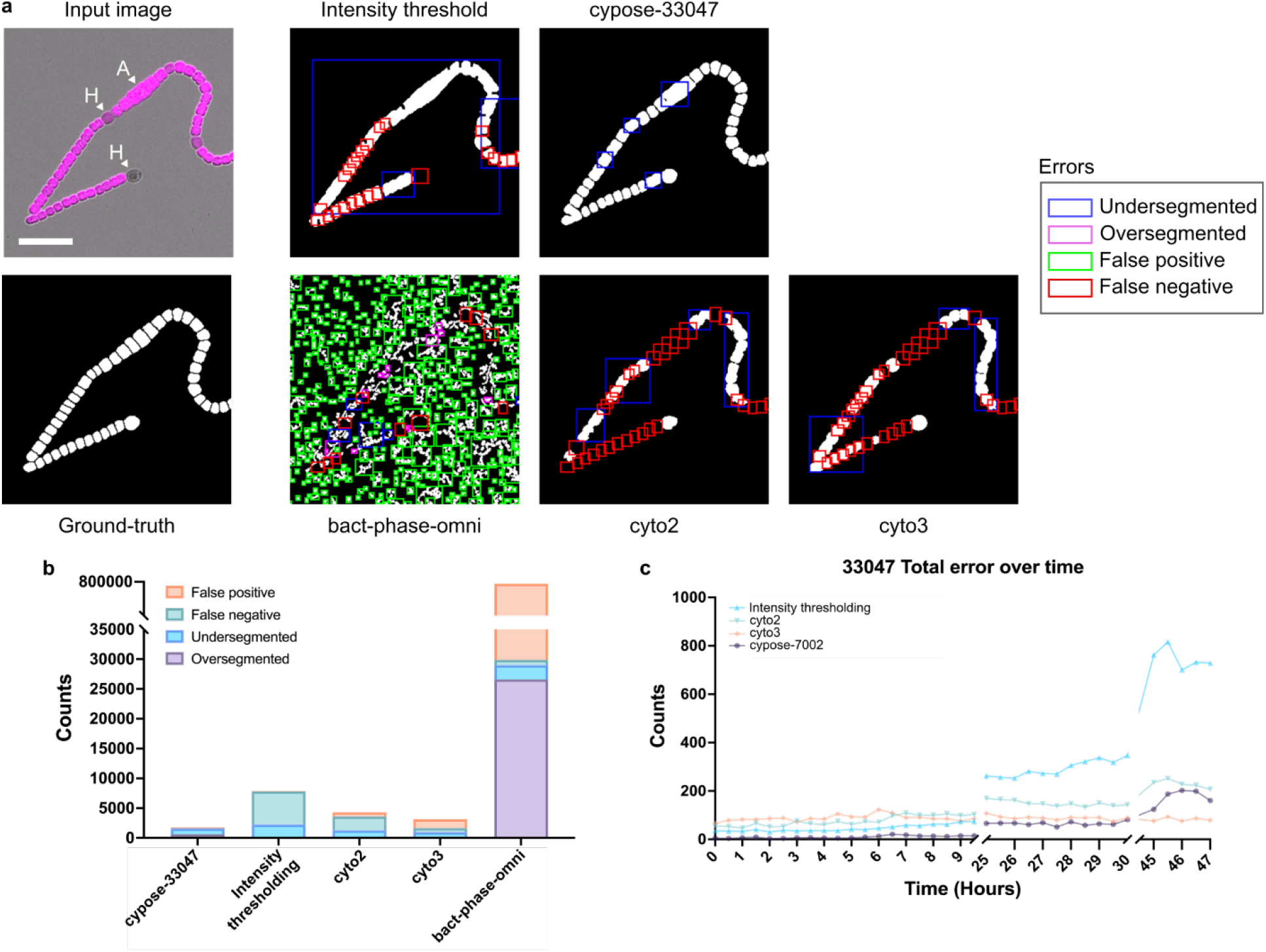
(a) Representative images of Anabaena 33047 showing the input image, ground-truth, and resulting masks from the intensity-thresholding algorithm and the various machine learning models. The input image is composed of two channels: brightfield (grayscale) and chlorophyll fluorescence (magenta). The heterocysts (labeled H) showed reduced or no chlorophyll fluorescence. An akinete cell is present in the image (labeled A). Scale bar indicates 20 µm. (b) Bar plot showing the total number of segmentation errors. (c) Plot showing the total number of errors at each frame of the benchmark movie. The line for bact-phase-omni was excluded for clarity (data shown in Fig. S4).

More detailed analysis of the errors showed that the majority (52%, Table 1) of errors in cypose-33047 were due to under-segmentation (Fig. 2b, S4). However, our model still showed better results when compared to the next best model, cyto3 (cyto3 had 688 under-segmentation errors compared to 81 for cypose-33047). We also found that our model was more accurate at segmenting akinetes than the other tested models and was resilient to other imaging artifacts (S5).

### Development of Cyclass to classify cellular phenotypes in a single image

Microscopy-based assays allows individual cellular phenotypes to be observed, for example in microbiome^22^ or competition assays^23,24^ or to probe population heterogeneity^8,25^. Typically, cellular phenotypes are identified in post-processing by filtering using physical properties, such as size or growth rate, or by measuring the intensity of labelling with different fluorophores. However, developing these computational filters can be challenging if the phenotypes are not significantly different from each other or if a phenotype is not easily described by a single parameter.

Here, we describe a method to train a convolutional neural network (CNN) based classifier, named Cyclass (Fig. 3a), to identify different cellular phenotypes in an image. By using a CNN, we were able to train a model to recognize different phenotypes directly from input images without the need for explicitly designing filters. We note that, unlike the Cypose segmentation models, classifier models are not easily generalizable to different imaging conditions (e.g., number of available channels) or different phenotypes. Thus, it is likely that new classifier models must be trained for different applications. The Cyclass framework provided in the codebase enables users to train their own models using images consisting of up to 6 different channels.

**Fig 3:**
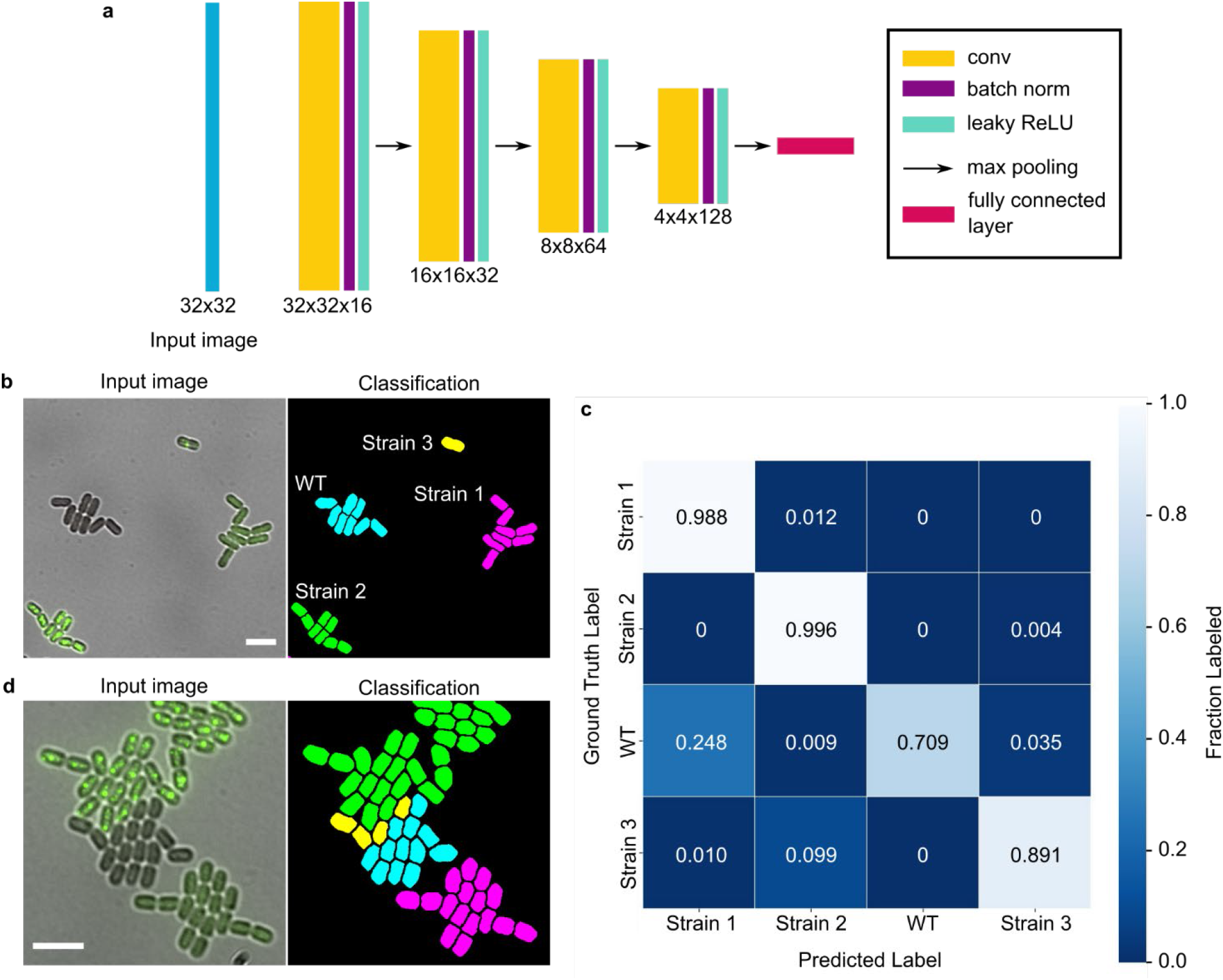
(a) Classification network architecture. The values provided indicate the size of the input image and the sizes of the feature maps in each layer. (b) Representative images showing the input image (only brightfield and one of the GFP channels are shown here for clarity; the full set of channels are shown in S7) and a recolored mask showing the classification of four strains of PCC 7002 cells. The strains have differently localized GFP: WT (no GFP labelling), Strain 1 (diffuse in cytoplasm), Strain 2 (carboxysome), and Strain 3 (procarboxysome). (c) Confusion matrix of the classification model. (d) Image showing the most common classification error, where WT (cyan) is misclassified as Strain 3 (yellow). These errors occur primarily when the colonies grow close together. The scale bars indicate 5 µm.

As proof-of-principle, we trained a mode, named cyclass-7002, to classify four cell types of co-cultured PCC 7002 mutants with differently localized GFP: Strain 1 had GFP freely diffused throughout the cytosol, Strain 2 had GFP localized to the carboxysome, Strain 3 had GFP localized to the procarboxysome, and the WT strain had no GFP. Details of each strain are available elsewhere^26^.

A representative image showing the result of cyclass-7002 is shown in Fig. 3b. To visualize the results, we used the classification values from the spreadsheet to color the cell mask. To quantify the accuracy of the classification, we calculated the overall IoU (0.919), precision (0.958), and recall (0.958) scores. The confusion matrix was also calculated and is shown in Fig. 3c. When analyzing the error, we found that they primarily occurred in cells bordering merging colonies or in cells which exhibited intermediate phenotypes (Fig. 3d). This is likely because the input image size of 32×32 is larger than a single cell, and information from neighboring cells could affect the resulting classification.

## Discussion and conclusions

In summary, we have developed machine learning models for segmentation (Cypose) and classification of single cyanobacterial cells (Cyclass) in imaging datasets. These models can be used independently or within a pipeline in our previously developed CyAn software^3^ (Fig. 4). We have shown that the fine-tuned segmentation models generated from the generalist Cellpose models can outperform the originals, even when using images of cells which were highly distinct (i.e., bacteria) compared to those used to train the generalist models (i.e., primarily mammalian cells). Compared to our previous traditional intensity-based thresholding approach, these new models enable segmentation of challenging situations, such as PCC 7002 in dense colonies, and for filamentous bacteria like ATCC 33047. The fine-tuned models were also able to segment a variety of cell morphologies, allowing for broad application to images of cyanobacteria with different phenotypes.

**Fig 4:**
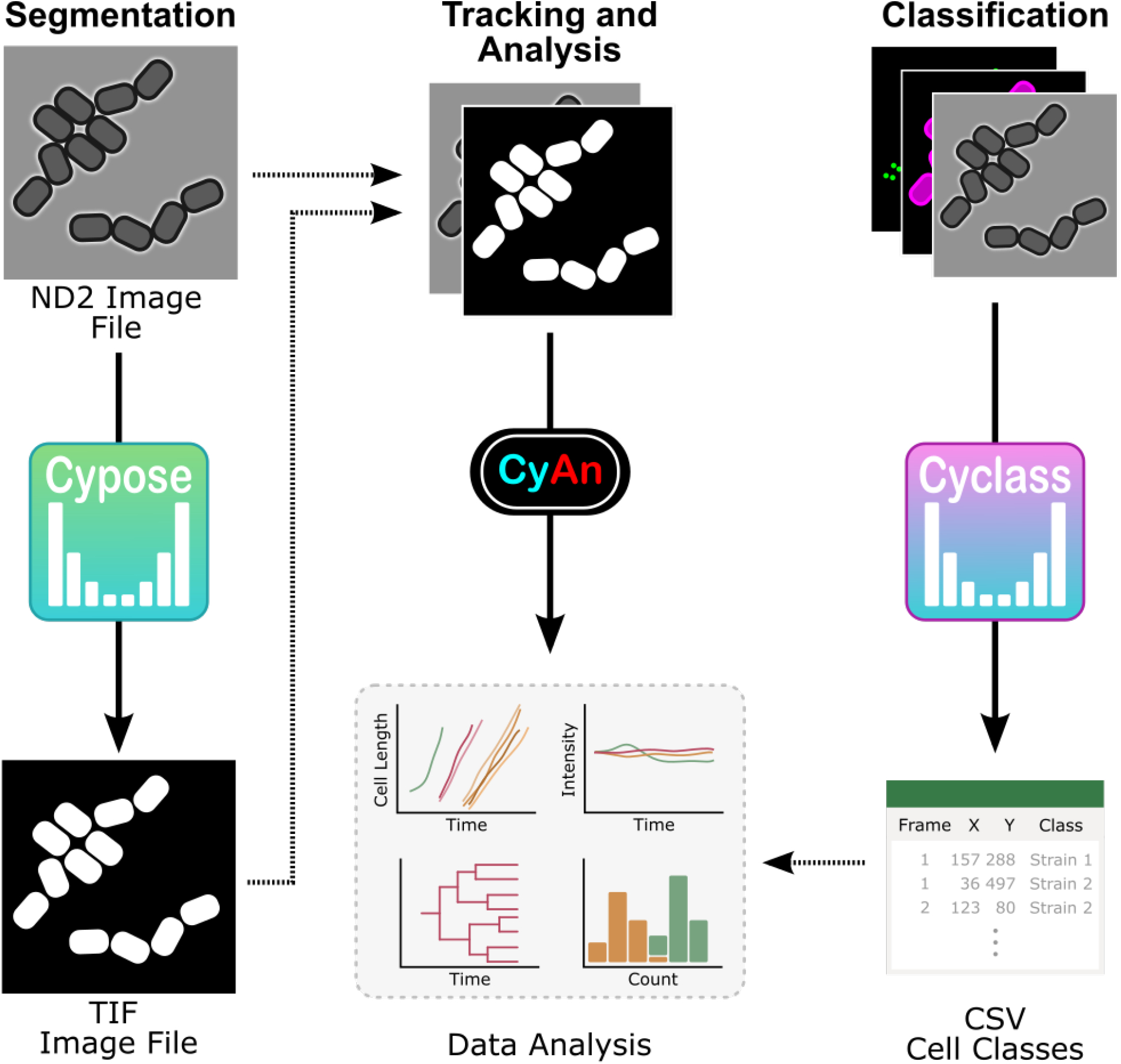
Overview of developed pipeline. Brightfield images are run through the appropriate Cypose model to generate cell masks, which are then exported as TIFF files. These masks, along with the original images, can then be input into our CyAn software package for tracking and data analysis. For images with different cell types, the images can also be run through a relevant Cyclass model to identify different cell classes. These classifications are saved in a CSV file and subsequently used to inform downstream data analysis.

Additionally, we investigated whether scratch-trained models performed better than fine-tuned models. We found that fine-tuned models trained from existing models provided the best balance between accuracy and resources required. Scratch-trained models offered little or no advantage over transfer learning.

Finally, we demonstrated that a cell classification model can be trained to classify different cellular phenotypes within a single image. Compared to traditional approaches, the cell classification model does not require the user to manually define filters (e.g., cell size or intensity) to identify different cell types, making analysis more robust and less prone to observer bias.

These new models advance the current state of image analysis for cyanobacterial imaging experiments by improving cell segmentation and classification. We believe that these models will enable future microscopy-based assays where different mutants or species are grown in the same conditions and imaged within a single field of view.

## Methods

### Dataset generation

#### Strain cultivation

To generate the training datasets, we acquired new images, as well as repurposed datasets from previous experiments, both reported and unreported. Details for all strains are provided in references cited in Table S1.

*Synechococcus* sp. PCC 7002 strains were cultivated in AL-41 L4 Environmental Chambers (Percival Scientific, Perry, IA) at 37°C under constant illumination (∼150 µmol photons m^−2^ s^−1^) by cool white, fluorescent lamps, under either ambient or elevated (3%) CO_2_ conditions. Cultures were grown in 25 ml of A+ media in orbital shaking baffled flasks (125 ml) contained with foam stoppers (Jaece Identi-Plug), or on pH 8.2 A+ media solidified with Bacto Agar (1%; w/v). Antibiotics were added for routine growth of strains (kanamycin, 100 µg/ml; gentamycin, 30 µg/ml), depending on the genotype.

*Anabaena* sp. ATCC 33047 was obtained from the Pakrasi lab and grown at 37°C in BG11 media. All preculturing occurring in 25 mL liquid cultures with 100 rpm orbital shaking in 125 ml baffled flasks with foam stoppers. Liquid and agar cultures were grown in an AL-41 L4 Environmental Chamber (Percival Scientific, Perry, IA) at 37°C under constant illumination at ∼150 μmol photons m^−2^ s^−1^ by cool white, fluorescent lamps in ambient air.

#### Microscopy

Images were taken using a Nikon TiE inverted wide-field microscope. Temperature and CO_2_ were controlled with an environmental chamber (Okolab) outfitted with a ProCO_2_ P120 Carbon Dioxide Single Chamber Controller (BioSpherix). Growth light was provided by a transilluminating red light emitting diode (LED) light source (Lida Light Engine, Lumencor). Fluorescence imaging was carried out using a highspeed light source (Spectra X Light Engine, Lumencor). NIS Elements software (version 5.11.00, 64-bits) with the Jobs acquisition upgrade was used to control the microscope. Image acquisition was performed using an ORCA-Flash4.0 V2+ Digital sCMOS camera (Hamamatsu) with a Nikon CF160 Plan Apochromat Lambda 100× oil immersion objective (1.45 NA) for PCC 7002 or Nikon CFI Plan Apochromat Lambda D 20× air objective (0.80 NA) for ATCC 33047.

To acquire time-lapse datasets, PCC 7002 cells in exponential or early linear phase were diluted to 0.14 OD_730_. 1 µL was spotted onto a 1% agarose A+ pad and allowed to dry (20 min). The pad was then inverted into an imaging dish, which was then wrapped in parafilm to keep the pad from drying out. The cells were preincubated at 37°C for 1 hour in the dark. Images were taken every 20 minutes. Cells were constantly illuminated with red light except during fluorescent imaging.

ATCC 33047 cells were grown in liquid culture to ∼1.00 OD_730_. 3-5 x 2 μL drops of cells were added to the imaging side of a 1% agarose w/v BG11 (or BG11-N to induce heterocyst differentiation) pad and allowed to dry. The pad was flipped on to an imaging dish (Ibidi μ-dish 35mm glass). The imaging dish was then sealed with parafilm and placed into the microscope. Images were captured every 20 or 30 minutes, depending on the movie.

### Data preparation

We used existing algorithms to assist with ground truth data generation. Initial cell masks were generated using either the CyAn Toolbox^3^ (version 1.3.4) running on MATLAB version R2020b or cyto2 running on Python3. These masks were then manually corrected using ImageJ/Fiji. Dead or overlapping cells were not annotated during this process.

### Model training details

#### cypose-7002

The fine-tuned PCC 7002 model was trained using the Cellpose v2 framework, starting with the pretrained cyto2 model. A training corpus of 6 movies, consisting of 413 frames with a total of 35,000 cells was used. We note that only the brightfield channel was used for training this model. Training was carried out on a single Nvidia A100 GPU using the pytorch framework. The final model was trained for 150 epochs. To benchmark the model, we used a separate movie which was never shown to the model during training. This benchmark movie consisted of 70 frames.

#### cypose-33047

As before, this model was trained using the Cellpose v2 framework, starting with the pretrained cyto2 model. The training dataset consisted of images of ATCC 33047 containing both brightfield and chlorophyll fluorescence channels. A training corpus of 4 movies, consisting of 233 frames and 68411 total cells was used. Training was carried out on a NVIDIA T40 GPU using pytorch. The final model was trained for 1250 epochs. To benchmark the model, a separate movie was used. This movie was cropped to a total of 36 representative frames showing different cell densities from the start, middle, and end of the full-length video. The ground-truth data was made using an early version of the cypose-33047 model, then manually corrected.

#### cypose-7002-scratch

This model was trained from scratch using the Cellpose v2 framework. Since a larger dataset is required to train a model from scratch, we used images from an additional 12 time-lapse movies in addition to the 6 movies used to train cypose-7002. The total training set comprised of 18 movies with a total of 2271 frames and approximately 125,040 cells. To account for the different cell morphologies, the training dataset included images from various PCC 7002 knockdown mutants (e.g., *ΔmurA, ΔftsZ*, and *Δftsh1-4*) which show morphological deviations from WT-cells (e.g. cell swelling, cell elongation). The cypose-7002 model was used to generate the initial mask. As before, the masks were then manually corrected to generate the final training dataset.

### Segmentation benchmarking

To validate our segmentation models, we calculated the pixel accuracy using the typical precision-recall metrics^16^. However, as discussed above, these metrics provide misleading statistics as errors such as over-segmentation tend to only involve a small number of pixels compared to the size of the cells. As an alternative, we developed an algorithm to identify and count specific segmentation errors. The main segmentation errors that are detected are: over-segmentation (when a predicted object is divided into more pieces than the ground truth), under-segmentation (when multiple objects are grouped together into a single large object), false negative (objects that were found in the ground truth, but are missing in the predicted masks), and false positive (objects which were found in the predicted masks, but are not in the ground truth). To avoid overcounting the number of errors, the algorithm allowed approximately 10% discrepancy between the ground truth and predicted masks.

### Training the Cyclass classification model

As shown in Fig. 3a, the Cyclass classification network architecture consists of a series of 4 sets of convolutional layers, with kernel sizes of 3×3. The feature map of each layer was batch normalized, and a leaky ReLU activation function was used. Each layer was followed by a 2×2 max-pooling layer with a stride of 2. A final fully connected layer was used for the classification task.

To train the model, we used images from a dataset consisting of 4 distinct cell genotypes/phenotypes, as described above (see also Supplementary Table 1). The input images consisted of 5 channels, including brightfield and four fluorescent labels (see S7). To generate a training set, we manually annotated images as belonging to one of the four cell types. The model was trained with an input image size of 32×32 pixels (about ∼1.5x – 2× cell size). It is interesting to note that during testing, smaller images appeared to perform worse suggesting that the network likely requires some information from neighboring cells. Conversely, input images that were larger might confuse the network as it includes too many other cells. The model was trained using the Adam algorithm, stopping at 60 epochs. To validate the model, a separate movie consisting of 70 frames was used as a benchmark.

The classifier model was integrated into an automated pipeline. To use the classifier, we first generated masks to identify individual cells. The masks were then used to obtain the centroid position of each cell. A region of 32×32 pixels around this location was then cropped from the original image. The cropped image was then fed into the Cyclass network which calculated class probabilities for the cell. The class corresponding to the highest probability was then output to a csv file and used in downstream analysis (Fig. 4).

## Acknowledgements

The authors would like to dedicate this paper to the memory of Prof. Jeff Cameron, who provided funding and served as a mentor throughout most of this work. We thank Dr. Hamadri Pakrasi for the ATCC 33047 strain. This work was supported in part by the Interdisciplinary Quantitative Biology (IQ Biology) program at the BioFrontiers Institute, University of Colorado, Boulder, the NIH/CU Molecular Biophysics Program, the NIH Biophysics Training Grant T32 GM145437, the National Institutes of Health under grant NIH AI156739, the National Science Foundation under grant MCB-2054085 and the Department of Energy under grants DOE DE-SC0018368, DE-SC0020361, and DE-SC0025606 to JCC.

## Author Contribution

CAH and ZLM conceived and designed the study. Sample preparation and data collection were performed by CAH, AA, and CB. CAH, AA, CB, and JWT prepared the ground-truth datasets. ZLM and CAH developed the code and performed initial training for all fine-tuned models. The final models were trained by ZLM (cypose-7002) and AA (all other models). JWT and JCC provided supervision of the project. The initial manuscript draft was written by CAH, ZLM, and JWT. CAH, ZLM, AA, CB, and JWT reviewed, edited, and approved the final manuscript.

## Data availability

The training datasets used for this study are available upon request.

## Code availability

All code and trained models can be downloaded from https://github.com/cameronlab/cypose.

