## Supplementary Information for "Machine Learning Models for Segmentation and Classification of Cyanobacterial Cells"

Supplemental Table 1. Description of the models used in this study, along with descriptions of the training datasets.

| Model name | Description | Training dataset | Reference |
| --- | --- | --- | --- |
| cyto2 | Cell cytoplasm segmentation model from Cellpose version 2.0. | Trained in two-channel images, where first channel is channel to segment and second is an optional nuclear channel | ^16^ |
| cyto3 | Cell cytoplasm segmentation model from Cellpose version 3.0. | 9 datasets; Trained to generate images which segment well from noisy images | ^18^ |
| bact-phase-omni | Bacterial segmentation model from Omnipose | Phase contrast images of assorted bacterial species with diverse morphologies and optical characteristics, 27,500 total cells | ^19^ |
| cypose-7002 | Fine-tuned model for segmentation of PCC 7002 cells | 6 movies of PCC 7002, 413 frames, 35,000 total cells | This study |
| cypose-7002-scratch | Scratch-trained model for segmentation of PCC 7002 cells | 18 movies of PCC 7002 WT, *∆murA, ∆ftsh1-4, ∆pdbH, ∆ftsZ,* 2271 frames, 125,040 total cells. | This study |
| cypose-33047 | Fine-tuned model for segmentation of filamentous Anabaena cells | 4 movies of ATCC 33047, 233 frames, 68411 total cells | This study |
| cyclass | Cell classification network | 9 movies of PCC 7002, 736 frames, 55,695 total cells | ^26^ |

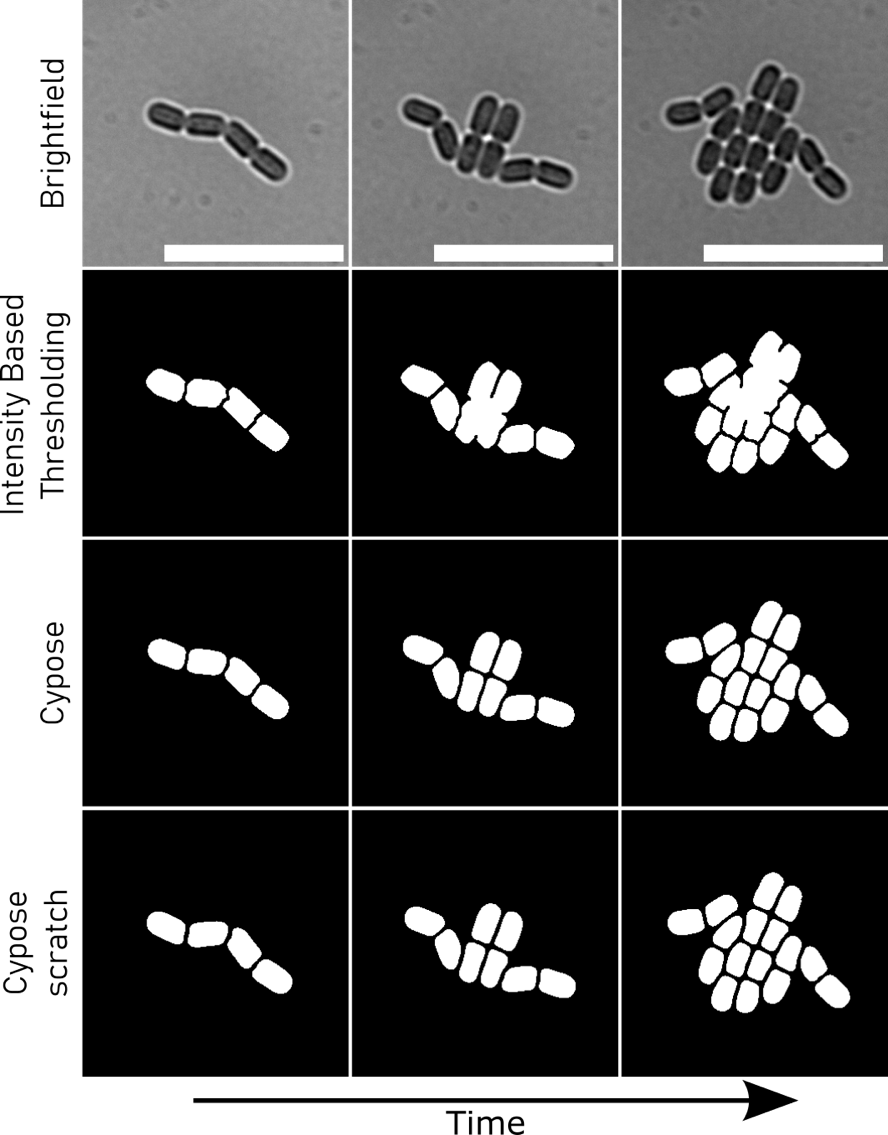

Supplemental Figure 1: Comparison of segmentation of 7002 between intensity-based thresholding, cypose-7002 and cypose-7002-scratch. Scale bars indicate 15 µm.

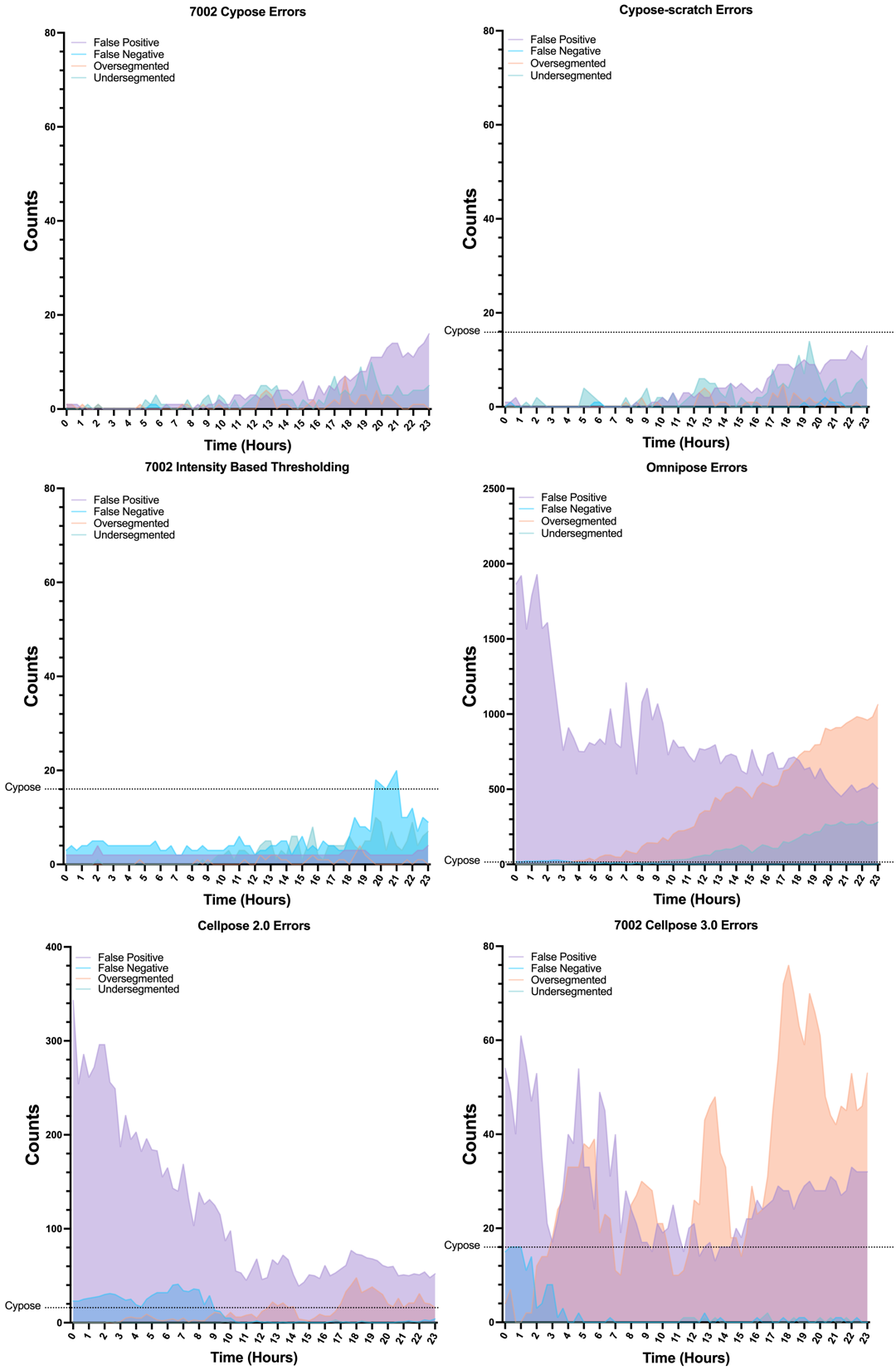

Supplemental Figure 2: 7002 segmentation error over time for all models

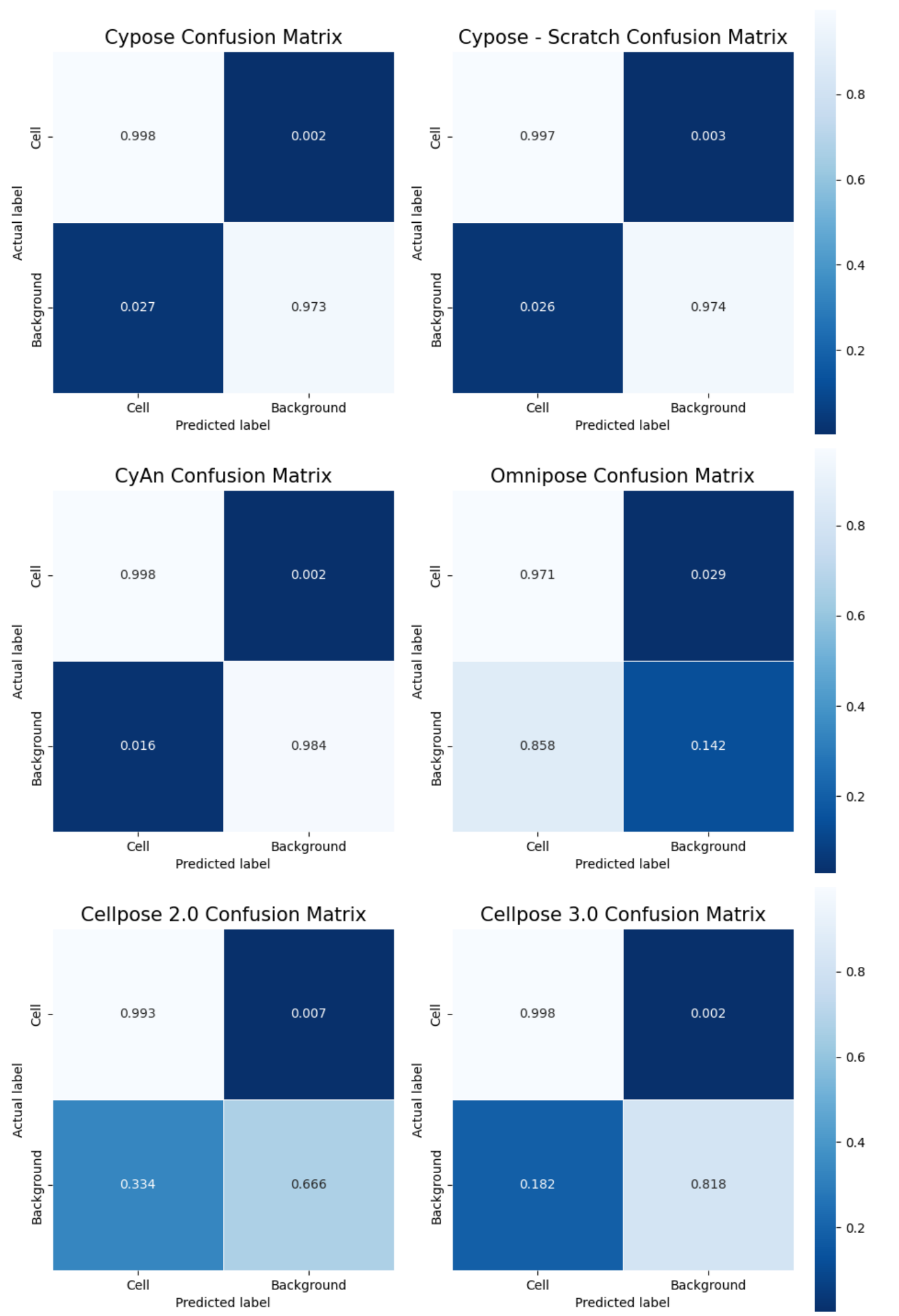

Supplemental Figure 3: Confusion matrixes for 7002 segmentation.

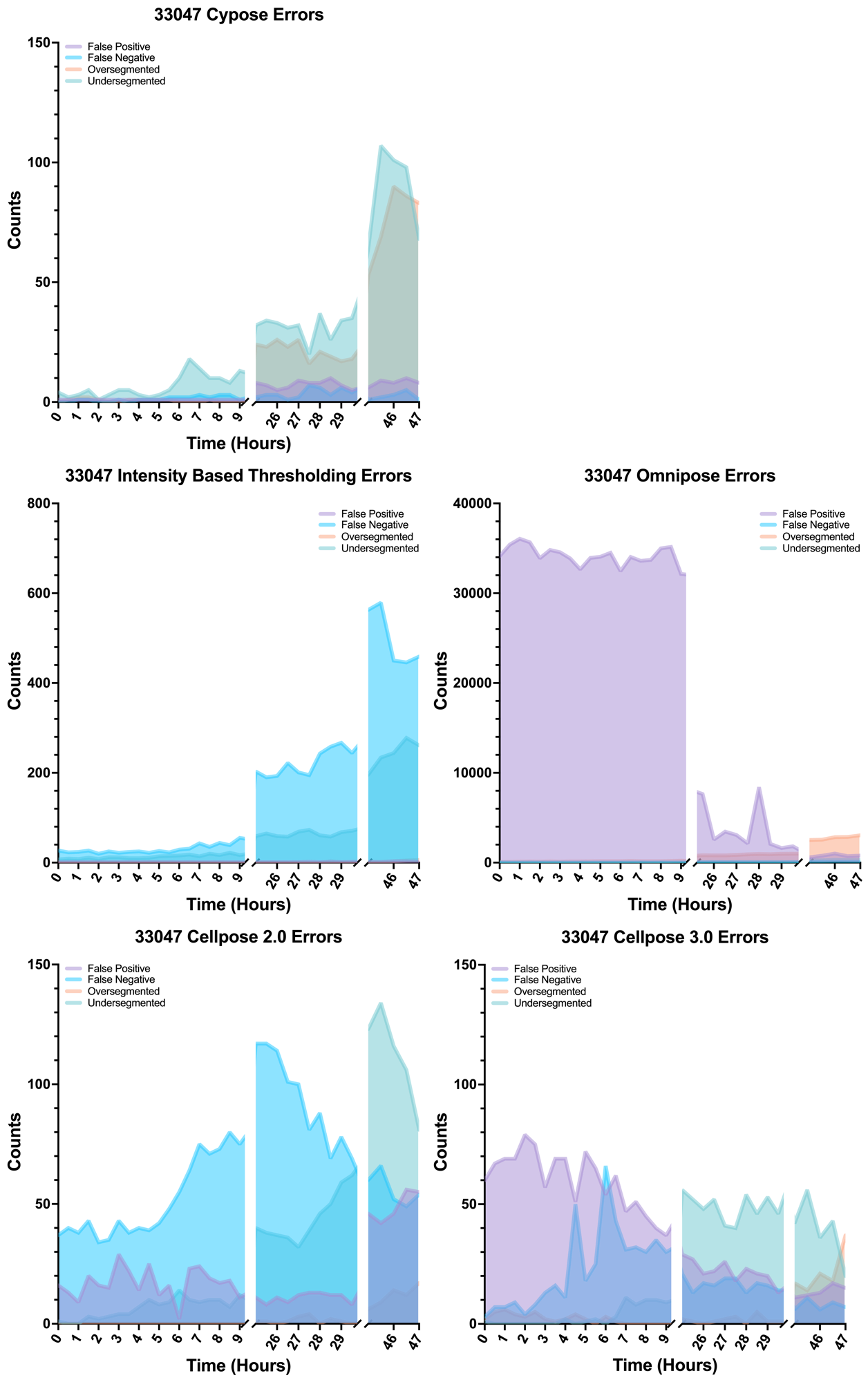

Supplemental Figure 4: 33047 segmentation errors over time for all models.

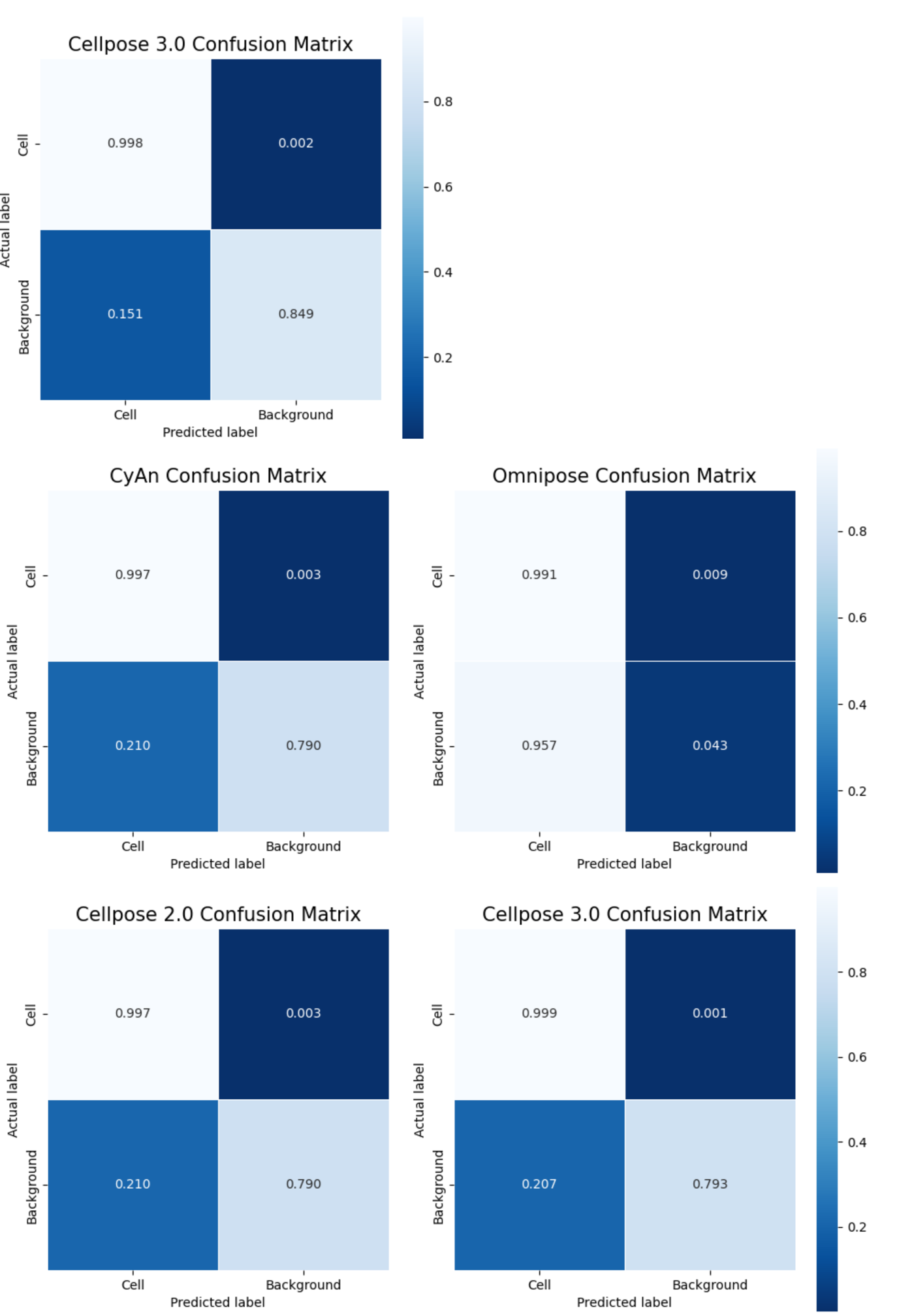

Supplemental Figure 5: Confusion matrixes for 33047 segmentation.

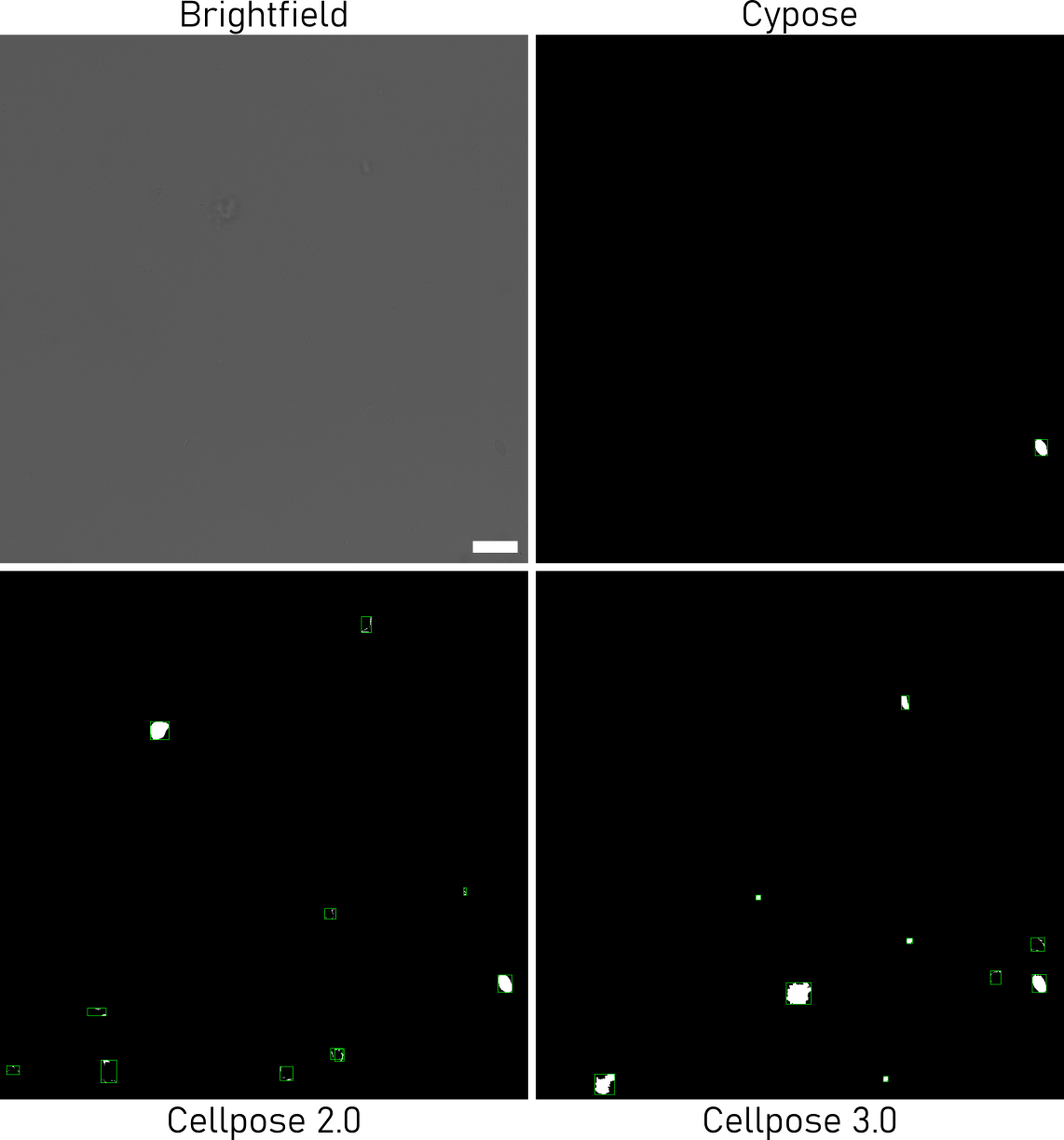

Supplemental Figure 6. 33047 segmentation comparisons showing false positive segmentation of debris and background.
